# Temporal coding expands the bandwidth of GPCR-mediated neuromodulation

**DOI:** 10.64898/2026.06.03.729927

**Authors:** Xinyi Jenny He, Mark von Zastrow

## Abstract

The advent of modern transcriptomic approaches has revealed a breadth of neuromodulatory G protein-coupled receptor (GPCR) co-expression far exceeding what was previously thought. This raises a fundamental question about neuromodulatory system architecture because there is a much smaller number of available G protein transducers. Does this impose an information ‘bottleneck’, or can a single transducer pathway distinguish the effects of different neuromodulators? We addressed this question by focusing on four GPCRs that are natively co-expressed in essentially all hippocampal pyramidal neurons and that signal through the canonical Gs-coupled cyclic AMP (cAMP) cascade. We show that each GPCR produces a similar cAMP elevation upon acute activation but the effects observed at downstream steps diverge, to the point that the GPCRs differ dramatically in their ability to drive a transcriptional response. We also show that they differ in their ability to maintain functional responsiveness, with neuropeptide receptors remaining responsive for hours but monoamine receptors rapidly losing responsiveness. We conclude that the effective chemical bandwidth of cellular neuromodulation by GPCRs is not limited by the number of available transducers, and that there exists additional temporal coding which enables individual neurons to distinguish a richer neuromodulatory input.

## Introduction

Neuromodulation is a critical form of chemical neurotransmission that is important for cellular and organismal adaptations to internal and environmental stimuli. These stimuli are conveyed through a vast chemical palette of ligands (which include monoamines, neuropeptides, and lipid metabolites) that are secreted and bind to modulatory receptors, the largest family of these being GPCRs. GPCR activation triggers a cascade of intracellular signaling events that can globally shape an animal’s brain state and behavior, from short term adaptations to long term encoding of memories. There are over 800 GPCRs encoded in the mammalian genome, most of which are expressed in the nervous system^1^. Despite this large repertoire of GPCRs, receptor signaling is mediated by only four G protein families. This discrepancy poses a fundamental bottleneck in the amount of neuromodulatory information the nervous system can communicate.

Traditionally, individual neurons or neuron classes were thought to express a limited subset of GPCRs, akin to the number of possible transducers. Therefore, the bottleneck might be solved at the level of single cell receptor expression, where neurons express a select few receptors that each couple to distinct G protein pathways to produce divergent effects. However, recent advances in single-cell transcriptomics reveal additional layers beyond this framework, with evidence that neurons may co-express up to 40 distinct GPCRs, far exceeding the classes of G protein transducers^2,3^. This implies that in any given neuron, there exists a multitude of GPCRs capable of coupling to the same transducer pathway, recapitulating the information bottleneck problem at the level of individual neurons.

This problem is particularly salient when considering the diverse neuromodulatory inputs available to a neuron. A clear example of this is the difference between the neuromodulatory systems of monoamines and neuropeptides. Monoamines and neuropeptides differ in their packaging, release, and clearance mechanisms which results in distinct ligand dynamics in the extracellular space^4–6^. However, surprisingly little is known about whether there are differences between monoamine and neuropeptide signaling pathways at the level of their cognate receptors. Recent observations point out that nearly all neurons have the capacity to express multiple GPCRs for both ligand classes^3,7^. This implies that individual neurons integrate signals from monoamine and neuropeptide GPCRs coupled to the same G protein, illustrating the bottleneck problem in a distinct physiological context. To address this problem, we set out to ask first, are GPCRs for monoamines and neuropeptides functionally expressed in the same neuron? If this is the case, are the responses produced by monoamine and neuropeptide GPCRs functionally distinguishable?

We investigate these questions by examining two monoamine GPCRs and two neuropeptide GPCRs that couple to the same signaling pathway. We discover functional co-expression and remarkable heterogeneity in signaling consequences in hippocampal neurons. These findings reveal that not all Gs-coupled GPCRs are created equal, and monoamine and neuropeptide GPCRs instead possess distinct signaling properties that determine the duration and functional consequences of their neuromodulatory responses.

## Results

### Functional co-expression of four native Gs-coupled GPCRs in primary hippocampal neurons

Based on a recent high-quality single-cell transcriptomic data set^2,3^, we predicted that individual glutamatergic neurons in the hippocampus co-express multiple GPCRs capable of coupling to the same G protein class. To test this prediction functionally, we focused on Gs-coupled GPCRs and assessed signaling through the canonical cyclic AMP (cAMP) / protein kinase A (PKA) cascade, utilizing a dissociated hippocampal culture preparation in which 90-95% of neurons are glutamatergic^8,9^. With the transcriptomic data as a guide, we selected two candidate monoamine receptors (β1-adrenergic receptor (β1AR) and D5 dopaminergic receptor (D5R)) and two candidate neuropeptide receptors (vasoactive intestinal peptide receptor 1 (VIPR1) and corticotropin-releasing hormone receptor 1 (CRHR1)) that are broadly expressed at the transcript level in nearly all excitatory neurons in the hippocampus (Figure 1A). Each of these GPCRs has been observed to increase excitability in pyramidal neurons in previous physiological studies^10–14^, but whether each GPCR initiates signaling in precisely the same neurons has not been determined.

**Figure 1.**
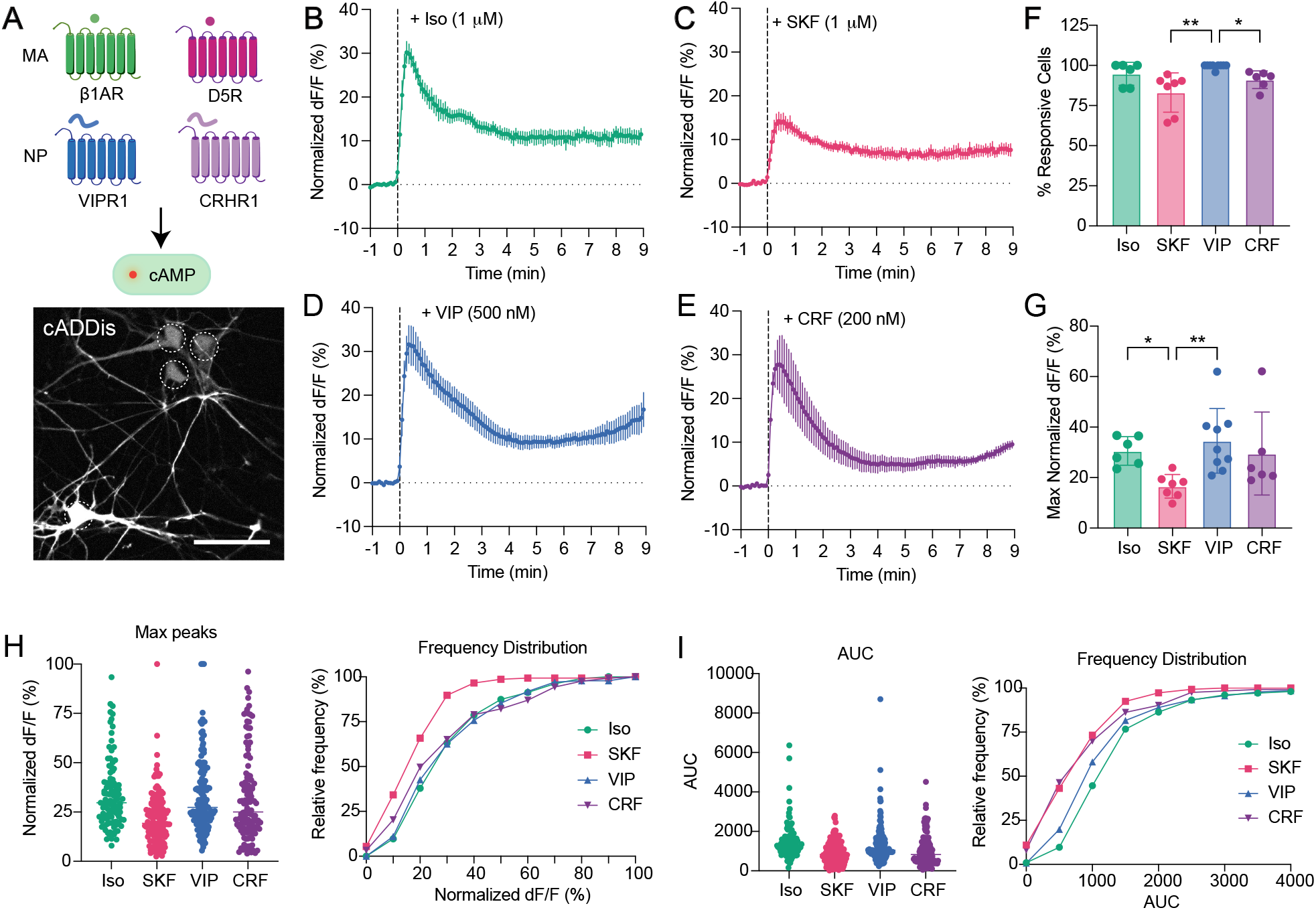
Global cAMP responses to application of selective agonists for each Gs-coupled GPCR. **A**. Schematic of four candidate Gs-coupled GPCRs – monoamine (MA) receptors β1AR and D5R and neuropeptide (NP) receptors VIPR1 and CRHR1 (top). Upon activation, Gs-coupled GPCRs canonically drive elevations in cAMP, which we measured using the fluorescent biosensor cADDis in primary hippocampal neurons (bottom). Dashed line ROIs indicate the soma that were measured for changes in fluorescence. Scale bar, 50 µm. **B**. The change in fluorescence upon addition of 1 uM isoproterenol (Iso), averaged across multiple neurons per dish (n = 6 dishes), normalized to 10 uM forskolin (FSK) added at the end of the time course (not shown). **C**. Normalized change in fluorescence with 1 uM SKF (n = 7 dishes). **D**. Normalized change in fluorescence with 500 nM VIP (n = 9 dishes). **E**. Normalized change in fluorescence with 200 nM CRF (n = 6 dishes). **F**. Percentage of imaged cells in a dish with a positive response to the agonist, defined as reaching 5% the FSK response within 1 minute of agonist application. Kruskal-Wallis test with Dunn’s multiple comparisons (*p < 0.05, **p < 0.005). **G**. The maximum change in fluorescence for the average response of a dish to each agonist, normalized to FSK. Kruskal-Wallis test with Dunn’s multiple comparisons (*p < 0.05, **p < 0.005). **H**. Peak responses for every individual neuron imaged across all imaging sessions, normalized to FSK (left). Frequency distributions of the max peaks, normalized to FSK (right). **I**. Area under the curve (AUC) for every individual neuron imaged across all imaging sessions (left). Frequency distributions of the AUCs (right).

We assessed functional signaling using well-characterized agonists capable of distinguishing between the selected candidates: SKF-81297 (SKF, 1 µM) for D5R, isoproterenol (Iso, 1 µM) for β1AR, vasoactive intestinal peptide (VIP, 500 nM) for VIPR1 and corticotropin releasing factor (CRF, 200 nM) for CRHR1. To test for the presence of functional GPCR expression, we measured the ability of each agonist to increase global cAMP concentrations in the cell body, using the real-time cAMP biosensor, cADDis^15^. If present, Gs-coupled receptors are expected to produce elevations in cAMP within minutes, therefore we monitored the effects of each agonist on cAMP for a 10 minute time course. At the end of the time course, we added the supra-physiological adenylyl cyclase activator forskolin (FSK, 10 µM) to maximally activate the sensor. This served both to normalize the agonist response and to verify that agonist responses were not saturating the sensor.

All of the agonists produced detectable cAMP elevation (Figure 1B-E) in the majority of cells imaged (Figure 1F). Iso produced responses in 95 ± 7% of cells, SKF produced responses in 83 ± 12% of cells, VIP produced responses in 99 ± 1% of cells, and CRF produced responses in 91 ± 6% of cells. This high degree of responsiveness indicated that the vast majority of neurons must co-express all of the candidate GPCRs tested. Further supporting this, we observed cAMP elevations to each of the agonists when applied sequentially to the same neurons (Figure S1).

We then compared the magnitude of the cAMP responses in two ways. First, we calculated the averaged responses across the cell population, measured at the peak. In this comparison, Iso, VIP, and CRF produced comparable cAMP responses while SKF produced a smaller response than VIP and Iso (Figure 1G). Second, we evaluated the discrete single cell responses produced by each agonist. While clear agonist responses were observed in nearly all neurons, the single-cell analysis revealed significant heterogeneity in individual responses. This was observed for the magnitude of individual peaks (Figure 1H) and the area under the curve (AUC) (Figure 1I). For the individual maximum peaks, the distribution of peaks generated by SKF was shifted to the left, indicating that the single-cell population of responses is skewed lower than the other 3 agonists (Figure 1H). Altogether, these results indicate that all 4 Gs-coupled receptors are functionally co-expressed, with β1AR, VIPR1, and CRHR1 producing comparable levels of global cAMP and D5R producing a less robust response.

### Differences in downstream PKA signaling between neuropeptide and monoamine receptors

To investigate if differences in response magnitude were evident further downstream in the canonical cAMP cascade, we next examined global PKA activity using the well-established PKA activity biosensor, ExRai-AKAR2^16^. Because increase in cAMP leads to increase in PKA activity, we expected to similarly see increase in biosensor fluorescence within minutes of bath application of the agonists, as measured at the cell body (Figure 2A). Once again, we applied FSK at the end of the time course to normalize agonist-driven activity and confirm that the biosensor was not saturated.

**Figure 2.**
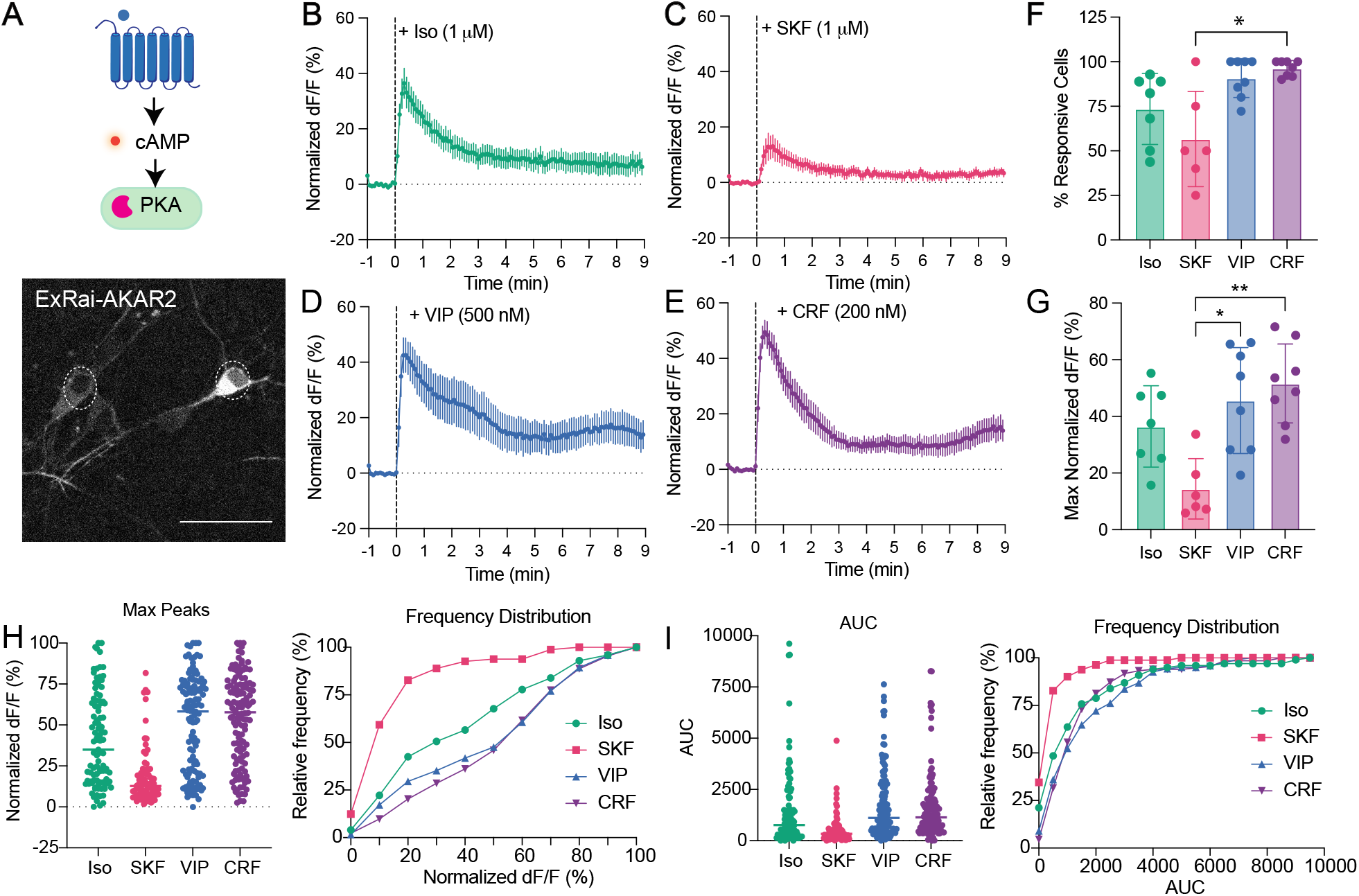
Global PKA responses to application of selective agonists for each Gs-coupled GPCR. **A**. Schematic of Gs-coupled GPCR signaling where activation of GPCR drives increase in cAMP concentration which then increases PKA activity (top). We measured PKA activity by expressing the green fluorescent biosensor ExRai-AKAR2 (bottom). Dashed line ROIs indicate the neuron cell bodies that were measured for changes in fluorescence. Scale bar, 50 µm. **B**. The change in fluorescence upon addition of 1 uM isoproterenol (Iso), averaged across multiple neurons per dish (n = 7 dishes), normalized to 10 uM forskolin (FSK) added at the end of the time course. **C**. Normalized change in fluorescence upon addition of 1 uM SKF (n = 6 dishes). **D**. Normalized change in fluorescence upon addition of 500 nM VIP (n = 8 dishes). **E**. Normalized change in fluorescence upon addition of 200 nM CRF (n = 8 dishes). **F**. Percentage of imaged cells in a dish with a positive response to the agonist, defined as reaching 5% the FSK response within 1 minute of agonist application. Kruskal-Wallis test with Dunn’s multiple comparisons (* p < 0.05). **G**. Maximum change in fluorescence for the average response of a dish to each agonist, normalized to FSK. Kruskal-Wallis test with Dunn’s multiple comparisons (* p < 0.05, ** p < 0.005). **H**. Peak responses for every individual neuron imaged across all imaging sessions, normalized to FSK (left). Frequency distributions of the max peaks, normalized to FSK (right). **I**. The area under the curve (AUC) for every individual neuron imaged across all imaging sessions, normalized to FSK (left). Frequency distributions of the AUCs, normalized to FSK (right).

Each agonist produced a rapid increase in global PKA activity with a similar time course as the upstream cAMP response (Figure 2B-E). Iso (74 ± 20%), VIP (91 ± 11%), and CRF (96 ± 4%) each produced a clear elevation of PKA activity in the vast majority of neurons imaged, whereas SKF did so in a smaller fraction of neurons (57 ± 27%) (Figure 2F). Furthermore, when averaged, the maximal PKA response produced by Iso, VIP, and CRF were comparable whereas the response produced by SKF was significantly lower (Figure 2G). When analyzed at the single-cell level, the global PKA activity varied considerably across individual neurons. On the individual neuron level, SKF application once again produced a distribution of peaks with smaller amplitudes (Figure 2H) and smaller AUCs (Figure 2I). Additionally, the distribution of peaks produced by Iso was shifted to the left of peaks produced by VIP and CRF, although distributions of AUCs were not different. These data indicate that the differences measured at the cAMP level are communicated downstream to distinctions in global PKA. Additionally, the degree of distinction between the receptors become more pronounced.

The next downstream step in the cascade is regulation of PKA activity in the nucleus, which is distinct from global PKA activity because it requires additional transport through nuclear pores. Therefore, we wanted to ask if differences among the co-expressed GPCRs in their ability to increase global PKA activity also manifest in the nucleus. To assess this, we expressed the nuclear-localized PKA activity sensor ExRai-AKAR2-NLS^17^ (Figure 3A). We measured fluorescence changes in individual nuclei upon bath-application of each selective agonist over a longer period (20 minutes) since changes in nuclear PKA are characteristically slower and more sustained than cytosolic PKA^17^.

**Figure 3.**
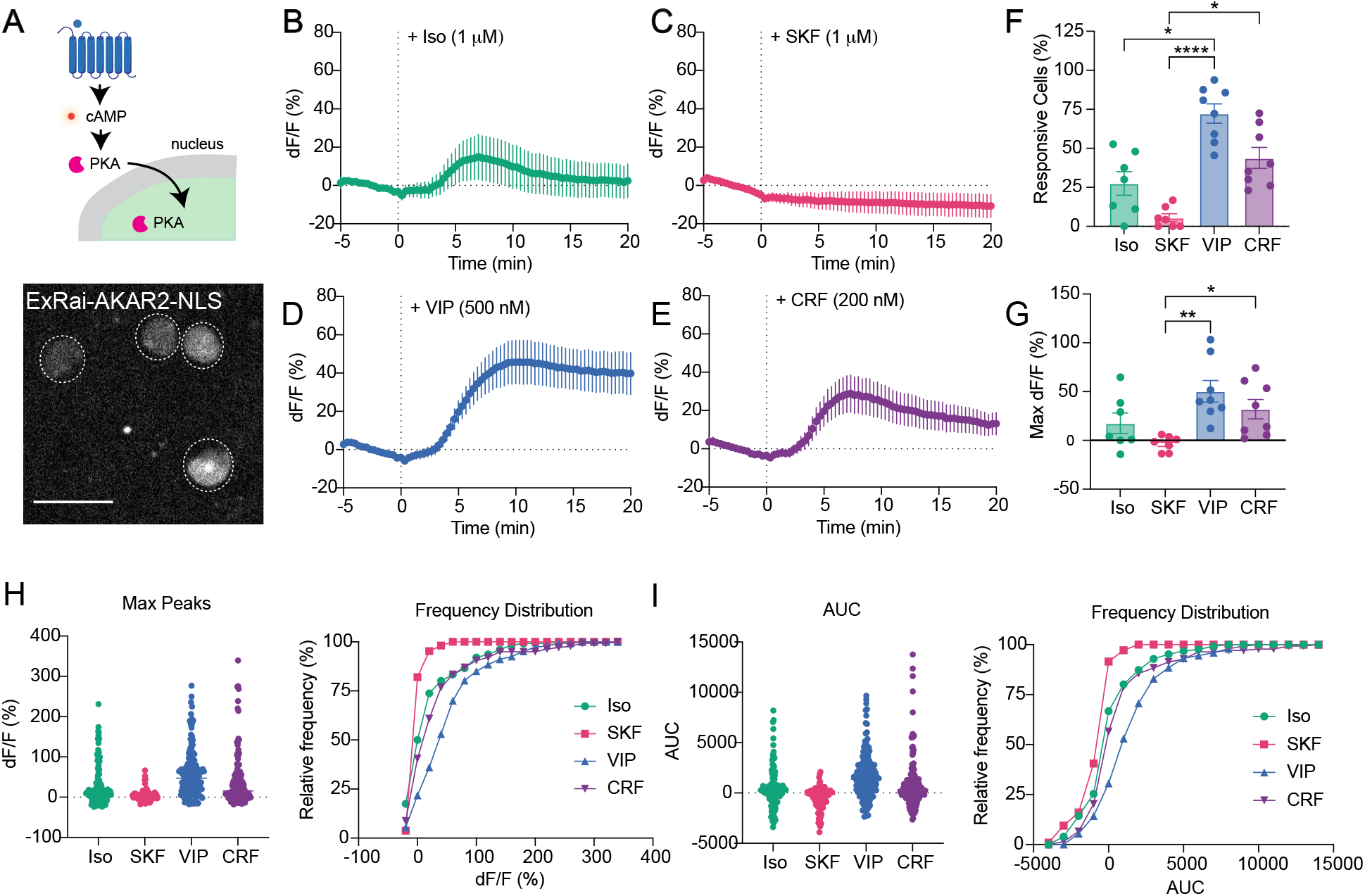
Nuclear PKA responses to application of selective agonists for each Gs-coupled GPCR. **A**. Schematic of Gs-coupled GPCR signaling where activation of GPCR drives increase in cAMP, then cytosolic PKA, which is transported to the nucleus to elevate nuclear PKA activity (top). We measured nuclear PKA activity by expressing the targeted green fluorescent biosensor ExRai-AKAR2-NLS which is restricted to the nucleus (bottom). Dashed line ROIs indicate 4 nuclei that were measured. Scale bar, 25 µm. **B**. The change in fluorescence upon addition of 1 uM isoproterenol (Iso), averaged across multiple neurons per dish (n = 7 dishes). **C**. Fluorescence change upon addition of 1 uM SKF (n = 7 dishes). **D**. Fluorescence change upon addition of 500 nM VIP (n = 8 dishes). **E**. Fluorescence change upon addition of 200 nM CRF (n = 8 dishes). **F**. Percentage of imaged cells in a dish with a positive response to the agonist, defined as reaching 20% change in fluorescence during the entire time course. Kruskal-Wallis test with Dunn’s multiple comparisons (* p < 0.05, **** p < 0.0001). **G**. Maximum change in fluorescence for the average response of a dish to each agonist. Kruskal-Wallis test with Dunn’s multiple comparisons (* p < 0.05, ** p < 0.005). **H**. Peak responses for every individual neuron imaged across all imaging sessions (left). Frequency distributions of the max peaks (right). **I**. The area under the curve (AUC) for every individual neuron imaged across all imaging sessions (left). Frequency distributions of the AUCs (right).

Agonist application produced variable changes in nuclear PKA activity that generally peaked between 5-10 minutes (Figure 3B-E). Iso produced clear elevations in activity in 28 ± 20% of cells while SKF only produced increases in 6 ± 6% of cells. Neuropeptides VIP and CRF produced marked increases in activity in 72 ± 18% and 44 ± 20% of cells, respectively. We again verified using bath application of FSK that the biosensor was not saturated at the nucleus by agonist application. However, because FSK-induced nuclear PKA activity continued to rise throughout the measured time course and did not reach a maximum steady state (Figure S2), the values reported in Figure 3 are not normalized to FSK.

We then compared the receptors by first examining the average fluorescence increases caused by each agonist across all measured neurons (Figure 3G). Here, SKF was inefficient at producing fluorescence increases in the nucleus (-3 ± 8%), similar to application of vehicle alone (-8 ± 13%) (Figure S2). VIP produced the greatest change in fluorescence (50 ± 31%), followed by CRF (32 ± 28%), and then Iso (18 ± 28%). Next, we quantified the single cell responses and once again we observed considerable heterogeneity in the maximum nuclear PKA elevation (Figure 3H) and the AUC (Figure 3I). Taken together, the differences observed followed the overall pattern as global PKA for SKF, Iso, and CRF. Additionally, we observed a relative enhancement in the ability for VIP to elevate nuclear PKA. This points to a divergence between the roles of distinct neuropeptide and monoamine receptors.

### Neuropeptide receptors drive sustained signaling with lasting cellular consequences

Having established heterogeneities in GPCR signaling at each step in the canonical cAMP-PKA cascade, we next assessed if these differences play a role in a functional consequence of PKA - regulation of gene expression through the transcription factor cyclic AMP responsive element binding protein (CREB). CREB-dependent transcription plays an important role in neuronal plasticity and survival and is known to be dependent on neuron activity^18–21^. To examine CREB induction, we developed a transcriptional reporter assay (CREB-GFP; UbC-mCherry) based on a previously published cAMP response element (CRE) reporter used in HEK293 cells^22^. Phosphorylation of endogenous CREB drives expression of eGFP downstream of a synthetic CRE motif which is introduced genetically into the neurons. Any constitutive expression of eGFP is suppressed by a dihydrofolate reductase (DHFR) degron, allowing for detection of transcription only during agonist treatment with the addition of degron inhibitor trimethoprim (TMP) (Figure 4A). We added a nuclear localized mCherry to visualize all neurons expressing the reporter.

**Figure 4.**
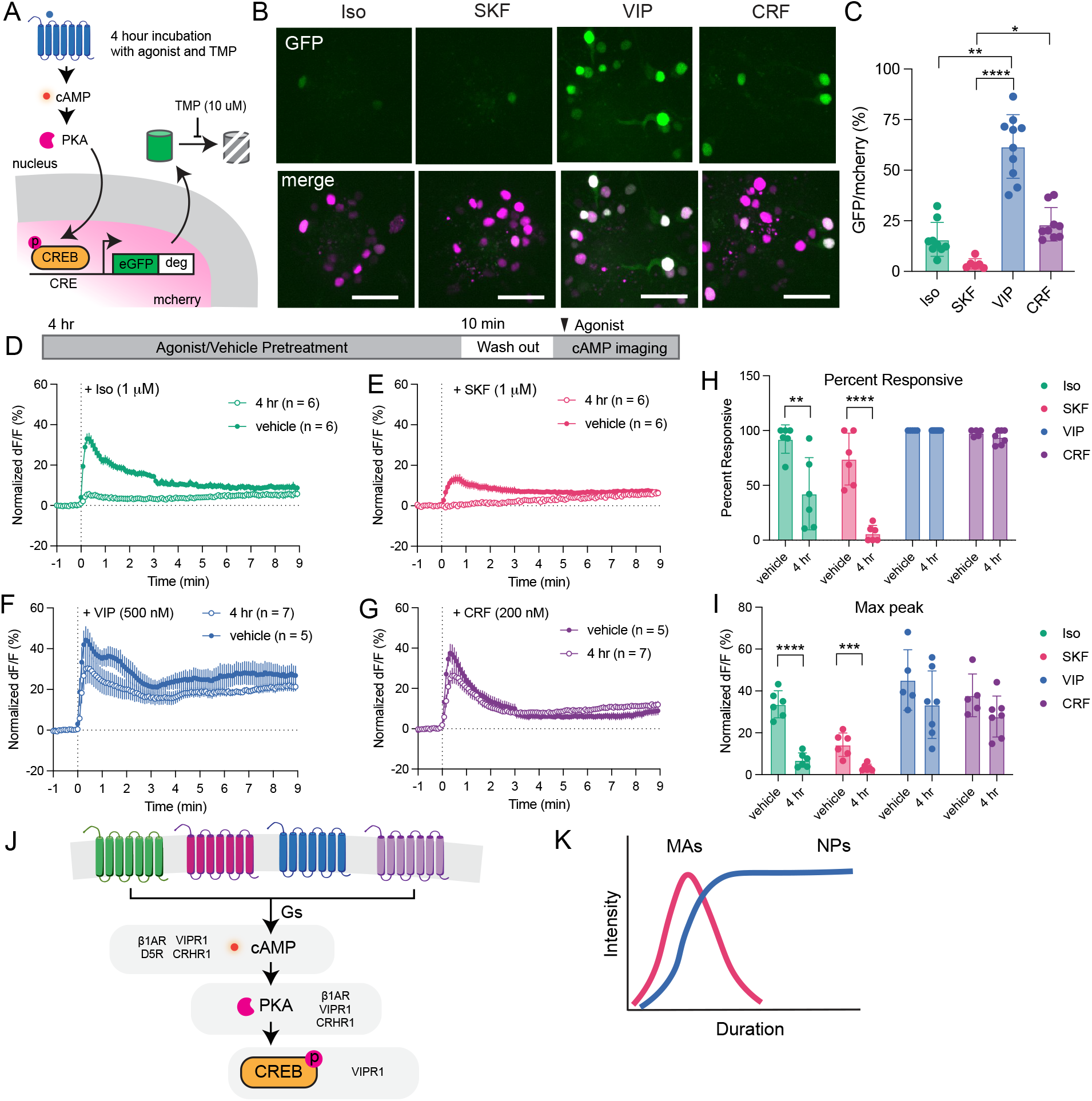
Long term functional consequences of selective activation of Gs-coupled GPCRs. **A**. Schematic of transcriptional reporter assay. Phosphorylation of CREB in the nucleus drives binding to synthetic CRE and transcription of eGFP. In the absence of trimethoprim (TMP), eGFP is degraded by degron. Nuclear mCherry is expressed within the same construct to label all cells transfected with the reporter. **B**. Example images of primary hippocampal neurons after 4 hours of incubation. GFP marks cells where transcription was turned on, while mCherry (magenta) marks all cells expressing the reporter. Scale bar, 50 µm. **C**. Percentage of reporter-expressing neurons that have induced GFP expression after 4 hours of drug treatment. Kruskal-Wallis test with Dunn’s multiple comparisons (* p = 0.027, ** p = 0.0013, **** p <0.0001). **D**. Schematic of experiment (top). Neurons expressing the cAMP sensor cADDis were incubated with either 1 uM iso or vehicle for 4 hours at 37 degrees. Neurons were then washed for 10 min and the fluorescence was measured upon addition of 1 uM Iso, normalized to 10 uM FSK applied at the end. **E**. Normalized fluorescence changes upon application of SKF after 4 hour pretreatment of SKF or vehicle. **F**. Normalized fluorescence changes upon application of VIP after 4 hour pretreatment of VIP or vehicle. **G**. Normalized fluorescence changes upon application of CRF after 4 hour pretreatment of CRF or vehicle. **H**. Percentage of neurons in a dish that respond to agonist following pretreatment by vehicle or the same agonist. Unpaired t test (** p = 0.0063, **** p < 0.0001). **I**. The averaged peak cAMP responses following pretreatment by vehicle or the same agonist. Unpaired t test (*** p = 0.001, **** p < 0.0001). **J**. Schematic summarizing the cAMP/PKA pathway and indicating the relative receptor effects at each stage of the pathway. **K**. Generalized model of monoamine (MA) neuropeptide (NP) signaling as a function of duration and intensity.

Incubation for 4 hours with TMP and adenylyl cyclase activator FSK induced GFP expression in 70 ± 11% of mCherry+ neurons, while TMP with vehicle treatment only induced GFP in 6 ± 3% of neurons (Figure S3). Incubating neurons for 4 hours with Gs-coupled receptor agonists evoked a wide range of transcriptional responses (Figure 4B). Notably, VIP produced a transcriptional response in 62 ± 16% of neurons, a greater proportion than that caused by incubation of SKF, Iso, and CRF (Figure 4C), and almost as great as FSK (Figure S3). SKF was remarkably weak in driving transcription (4 ± 3% of neurons), similar to the vehicle condition, and Iso and CRF produced transcription in a sparse subset of neurons (16 ± 8% and 23 ± 8%, respectively). As expected, the transcriptional response was dependent both on PKA activity and neuron activity as it was attenuated by the PKA inhibitor H89 (10 µM) and the Na^+^ channel inhibitor tetrodotoxin (TTX, 500 nM) (Figure S4). We verified the use of this reporter by staining for the induction of the endogenous immediate early gene *cfos*. One hour of agonist treatment produced the same trends in *cfos* activation by each of the agonists as with the CREB reporter (Figure S5).

The differences in transcriptional activation between GPCRs were remarkably large and compounded compared to the more nuanced distinctions upstream in the cAMP/PKA pathway. Therefore, we considered if there were additional disparities such as in the ability of the GPCRs to sustain long-term responsiveness to their cognate neuromodulator. To assess this, we incubated neurons with each agonist for 4 hours, washed out the agonist, and then measured the ability for the GPCRs to elevate cAMP (measured using cADDis) in response to a second agonist challenge (Figure 4D). We found that after 4 hours of pretreatment, Iso and SKF were ineffective at driving cAMP (Figure 4D and E). The percentage of responding cells and the peak cAMP response were reduced for Iso and SKF after pretreatment with agonist compared to vehicle (Figure 4H and I). On the other hand, after 4 hours of pretreatment with the neuropeptides VIP and CRF, neurons were still able to respond to agonist reapplication (Figure 4F and G). There was no difference between the percentage of responding cells and the peak cAMP response between vehicle pretreatment and agonist pretreatment for VIP and CRH (Figure 4H and I). Significant differences for monoamine receptors were even observed after only 10 minutes of Iso and SKF pretreatment, while neuropeptide receptor responses remained unchanged after 10 minutes of VIP and CRF pretreatment (Figure S6), demonstrating that these distinctions can be observed at more physiological time scales. Altogether, these results indicate that natively co-expressed GPCRs signaling through the shared cAMP/PKA cascade can produce heterogeneous neuromodulatory effects.

## Discussion

Current large scale multi-omics data sets are leading the field to understand that the complexity of neuromodulator ligand-receptor interactions is much greater than previously recognized. To fully appreciate the nuances in neuromodulatory communication, we must delineate the breadth of signaling achievable by GPCRs, first by determining if functional differences between receptors are even observable. Through a systematic examination of four Gs-coupled neuromodulatory GPCRs, we determined that multiple GPCRs are natively co-expressed in hippocampal neurons, and these GPCRs diverge in their long-term functions, with neuropeptide receptors better able to sustain downstream signaling than monoamine receptors (Figure 4J).

While there is scattered evidence of signaling differences between isolated neuropeptide and monoamine receptors, to our knowledge, the present results are the first to establish functional differences between native GPCRs within the same neuron. This is particularly timely, as emerging tools in neuroscience now allow direct assessment of neuromodulator release in awake, behaving animals with the development of monoamine and neuropeptide sensors^23–26^. As our insights into neuromodulator release expand, the receptor properties must be taken into consideration to infer the biological outcomes of neuromodulation. Our data demonstrate that GPCRs coupling to the same G protein pathway are not interchangeable. While we cannot speak for every neuropeptide and monoamine receptor in all neuron types, we can begin to draw some principles between the receptors for these two ligand classes.

Neuromodulation by these four receptors has been shown to have physiological effects in the hippocampus, but primarily for monoamines^27^. β-adrenergic receptors were initially described to increase the excitability of pyramidal cells via PKA^10,28^. Since then, they have been described to play a role in synaptic plasticity^29^ and contextual memories^30,31^. More recently, dopamine has been shown to play a role in spatial learning and fear conditioning^32–35^. CRH has been shown to be released in response to acute stress, impacting synaptic plasticity and regulating transcription^36,37^. VIP has been shown to have effects on pyramidal cell excitability^12^ and synaptic plasticity^38^. However, despite the presence of the VIP peptide in hippocampal interneurons, it is unknown under what conditions VIP is released to impact the local circuit. Taken together with our data, monoamine transmission may be primed to happen more frequently and transiently. Neuropeptide release through dense core vesicles, which requires more sustained stimuli and thus happens more rarely^4,5,39^, may be poised to enact broader and longer lasting changes to the neural circuit. More research on the stimuli evoking neuropeptide release and the timing and duration of release will be needed to fully understand the roles of neuropeptide signaling.

One of the most striking outcomes of this study is the heterogeneity in the ability for the four different receptors to induce transcription, with VIPR1 as the superior transcriptional activator. The exact mechanism behind VIPR1’s unique action is still to be determined, but many possibilities exist including subcellular location of signaling, strength of G protein coupling, and differences in non-cAMP effectors. We note that in HEK293 cells, VIPR1 signals strongly from endosome^40,41^, but it remains to be determined if this explains its strong transcriptional effect in neurons. The profound desensitization of monoamine receptors is a partial explanation for why β1AR and D5R are weak at driving long term transcription, however CRHR1 remains sensitive after 4 hours akin to VIPR1 (Figures 4F-I) yet drives a relatively modest transcriptional response. VIPR1 produces more nuclear PKA activity than CRHR1 (Figure 3D-G), which may compound into stronger transcriptional activation after 4 hours of cumulative signaling, suggesting another possibility. Regardless of the precise mechanism(s) underlying this difference, the present results indicate that transcriptional control elicited through natively co-expressed Gs-coupled GPCRs is remarkably receptor-specific.

From a global view of neuromodulation, our results pose an interesting juxtaposition – multiple Gs-coupled GPCRs are co-expressed but they can produce different downstream effects. This suggests that a single transducer pathway can communicate multidimensional information. The biochemical information produced by GPCR signaling emerges as a product of intensity and duration (Figure 4K). While short-term intensity of signaling by monoamine and neuropeptides is not dramatically different, neuropeptides can produce longer durations of signaling which can translate into profound differences in downstream processes such as regulation of gene expression. These fundamental principles provide another solution to the bottleneck problem, with neurons potentially able to decode signals based on duration of signaling. These coding principles can be overlaid on top of the neuromodulatory map to inform predictions about neuromodulatory systems and circuits.

## Materials and Methods

### Key Resources Table

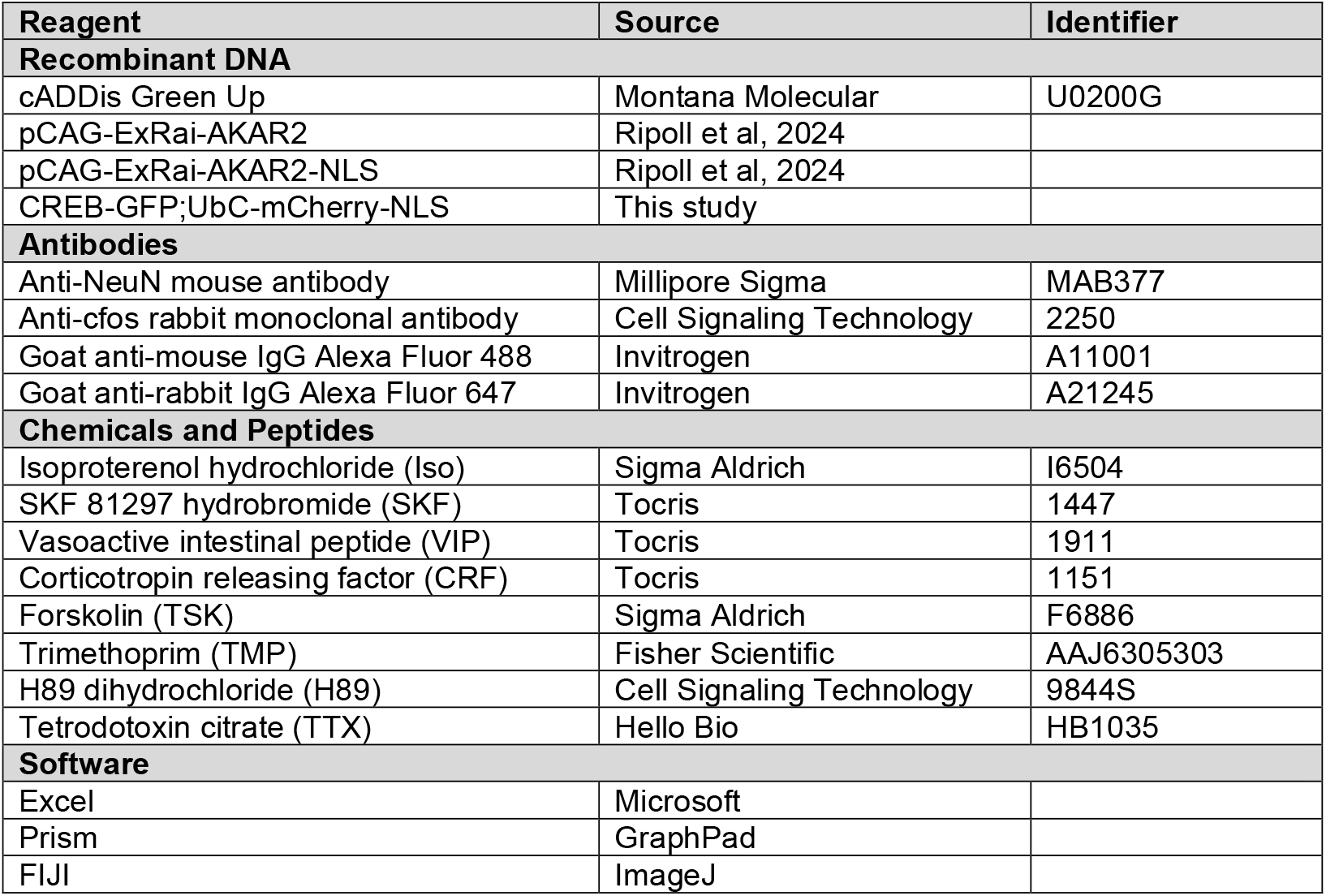

### Primary neuron culture

Primary hippocampal neurons were prepared from embryonic day 18 Sprague-Dawley rats (Charles River) dissected from a timed-pregnant female following euthanasia as previously reported. All procedures were performed according to the National Institutes of Health Guide for Care and Use of Laboratory Animals and approved by the University of California San Francisco Institutional Animal Care and Use Committee. The hippocampus was identified using a stereomicroscope and dissected into ice-cold Hank’s buffered saline solution calcium/magnesium/phenol red free (Thermo Fisher Scientific cat# 14-175-145). Hippocampi were dissociated in 0.05% trypsin/EDTA (University of California San Francisco, Cell Culture Facility) for 15 min at 37°C and washed 2x in Dulbecco’s modified Eagle’s medium (DMEM, Gibco) supplemented with 10% fetal bovine serum (FBS, University of California, San Francisco, Cell Culture Facility) and 30 mM HEPES (Gibco), then mechanically separated by trituration through flame-polished Pasteur pipettes. For electroporation, neurons were transfected using manufacturer’s instructions (Rat Neuron Nucleofector Kit, Lonza). Briefly, 6×10^6^ neurons were electroporated with up to 4 µg total DNA and 2×10^5^ neurons were plated onto 35 mm glass bottom dishes (Mattek) coated with poly-D-lysine (Sigma) in DMEM supplemented with 10% FBS. On day *in vitro* 4 (DIV4), media was exchanged for phenol red free Neurobasal medium (Gibco) supplemented with Glutamax 1X (Gibco) and B27 1X (Gibco). On DIV11, half of the cultured medium was exchanged with fresh, equilibriated medium and 2 µM cytosine arabinosine (Sigma-Aldrich) was added. All neurons were cultured in a humidified incubator with 5% CO_2_ at 37°C throughout, and imaged between DIV11 and DIV15. All experiments were performed with at least 3 independent cultures.

### cDNA constructs

To generate CREB-GFP; UbC-mCherry, we modified Cre-GFP reporter that was a gift from Willow Coyote-Maestas^22^. A gene fragment containing UbC-mCherry-3xNLS was synthesized by Twist Bioscience and inserted into the PCR-amplified Cre-GFP reporter between the attB site and the poly(A) signal with In-Fusion cloning (Takara Bio 638946).

### Live fluorescence imaging and quantification

For the cAMP assay, neurons were transduced with the Green Up cAMP biosensor (Montana Molecular) according to the manufacturer’s instructions. Live imaging was performed in HEPES buffered saline solution (HBS) adjusted to pH 7.4 containing, in mM: NaCl 120, KCl 2, CaCl 2 2, MgCl 2 2, Glucose 5, HEPES 10 and osmolarity was adjusted to 270mOsm. Immediately prior to imaging, neurons DIV 11-15 expressing biosensor were washed 1x with HBS and media was replaced with HBS. Live imaging was carried out on an inverted Nikon Ti2-E microscope with a CREST X-Light V2 large field of view spinning disk confocal using a 40x oil objective.

For cADDis and ExRai-AKAR2 time courses, images were taken every 5 s with a 488 nm laser for a total of 144 images per time course. Selective agonists Iso (1 µM), SKF 81297 (1 µM), VIP (500 nM), and CRF (200 nM) were added by bath application after a 1 min baseline. Forskolin (FSK, 10 μM) was added at 10 minutes for the last 2 minutes of the time course. For ExRai-AKAR2-NLS time courses, images were taken every 20 s with a 488 nm laser for a total of 76 images per time course. Agonists were added after a 5 minute baseline, and no FSK was applied.

Image processing and quantification was performed using FIJI Software (ImageJ). First, images were registered to correct for movement using the built-in FIJI plug-in StackReg, Translation. For cADDis and ExRai-AKAR2 images, background was subtracted using the built-in function Subtract Background, Rolling ball, 50. Then, ROIs were drawn around individual cell bodies and the mean fluorescence in the ROI was extracted for each cell throughout the time course. The fluorescence values per cell were baseline subtracted to calculate dF/F, then normalized per cell to its FSK max as 100%. The average time course for all neurons within a dish was calculated, as well as the time courses for each individual cell. Positive responses were defined as having responses greater than 5% of FSK within the first minute after agonist addition. For ExRai-AKAR2-NLS images, a uniform background subtraction was applied according to the lowest gray values of the image. Then, ROIs were drawn around individual nuclei and the mean fluorescence in the ROI was extracted for each nuclei throughout the time course. The fluorescence values per cell were baseline subtracted to get dF/F, and the average time course for all neurons within a dish was calculated. Positive responding cells were defined as having responses greater than 20% dF/F over the entire time course. Note that the cutoff value is different because these values were not normalized to FSK.

For agonist rechallenge experiments, neurons expressing cADDis were pretreated with agonist or vehicle for 4 hours at 37°C. Prior to imaging, neurons were washed 2x with HBS and the media was replaced with fresh HBS for 10 minutes prior to imaging of agonist rechallenge. Imaging and quantification was performed the same as described for cADDis above.

### Transcription reporter assay

For the transcription reporter assay, neurons DIV 11-15 expressing CREB-GFP-mCherry were treated with agonists and trimethoprim (TMP, 10 µM) in neurobasal for 4 hours at 37°C. Prior to imaging, the media was replaced with prewarmed HBS. Imaging was carried out on the CREST X-Light V2 large field of view spinning disk confocal with a 20x air objective. Z-stacks were taken at 1 µm steps using a 488 nm laser for GFP and 546 nm laser for mCherry, for a total of three 900 x 900 µm^2^ images per dish. Images were quantified in FIJI to extract cell counts using a custom macro to automate the process. Briefly, a maximum projection was created for each channel and the background was subtracted using a rolling ball subtraction. Then, the set threshold and analyze particles tools were used to extract cell outlines and numbers. Thresholds and particle sizes were selected by choosing values that detected most positive cells without falsely identifying noise, and values were kept consistent for each experiment. From the individual channel cell counts, the percentage of GFP cells compared to mCherry cells was quantified per image, and the average percentage for 3 images in a dish is represented as a replicate.

### Immunostaining

For the labeling of endogenous *cfos*, neurons DIV 14 were treated with FSK (10 µM), vehicle, or agonists for 1 hour prior to wash with phosphate-buffered saline (PBS, Life Technologies cat# 14040216) and fixation for 10 min at room temperature in 4% paraformaldehyde/PBS with sucrose (Fisher Scientific cat# AAJ19943K2). Neurons were blocked in 0.2% Triton X-100 and 3% bovine serum albumin (Sigma-Aldrich, A7030) in PBS for 1 hr at RT. Primary antibodies *cfos* (1:1000) and NeuN (1:1000) were diluted in blocking solution and incubated overnight at 4°C. Neurons were then washed three times in 0.2% Triton X-100 in PBS, 5 min, at RT. Secondary antibodies were diluted in blocking solution and labeling was performed at RT for 1 hr. Neurons were washed one time, 5 min with 0.2% Triton X-100 in PBS, then two more times with PBS, and stored in 1x PBS. Neurons were imaged by confocal microscopy using the CREST X-Light V2 large field of view spinning disk confocal with a 20x air objective. As for the transcription reporter assay, 3 z-stack images were taken per dish using a 488 nm laser and a 647 nm laser. Images were acquired and quantified using the same procedure as described above for the transcription reporter assay, except using a percentage of *cfos* cells / NeuN expressing cells.

## Statistics

Quantification of data in the text are presented as mean ± standard deviation (SD). In the figures, all data are presented as mean with error bars representing standard error of the mean (SEM). All experiments were carried out on at least 3 independent neuronal cultures, with the precise sample size indicated in the figure legends. Statistical analysis was performed using Prism 11 (GraphPad Software). For multiple comparisons, Kruskal-Wallis test with Dunn’s multiple comparisons was used for all except the agonist re-challenge experiment, where unpaired t tests were used.

## Acknowledgements

We thank Matthew Howard and Willow Coyote-Maestas for providing original CREB reporter construct. We thank Lea Ripoll, Rita Fagan, Emily Blythe, and Nicole Fisher for expression constructs and cloning assistance. We thank all members of the von Zastrow lab for valuable discussion and feedback on the manuscript. Imaging experiments were carried out at the Center for Advanced Light Microscopy at UCSF on CREST/C2 Confocal obtained using grants from the UCSF Program for Breakthrough Biomedical Research funded in part by the Sandler Foundation and the UCSF Research Resource Fund Award. We thank Nico Sturrman, Kari Herrington, and Micaela Lasser for their technical support and expertise. This work was supported by the National Institutes of Health National Institute on Drug Abuse (R01DA010711 and R01DA012864 to M.v.Z.). X.J.H. was supported by an NIH/NRSA BRAIN Initiative Postdoctoral Fellowship (F32MH136662).

## Author Contributions

X.J.H. and M.v.Z. conceived of the study and designed experiments. X.J.H. performed and analyzed experiments. X.J.H. and M.v.Z. wrote the paper.

