## Supplemental Figures for "Temporal coding expands the bandwidth of GPCR-mediated neuromodulation"

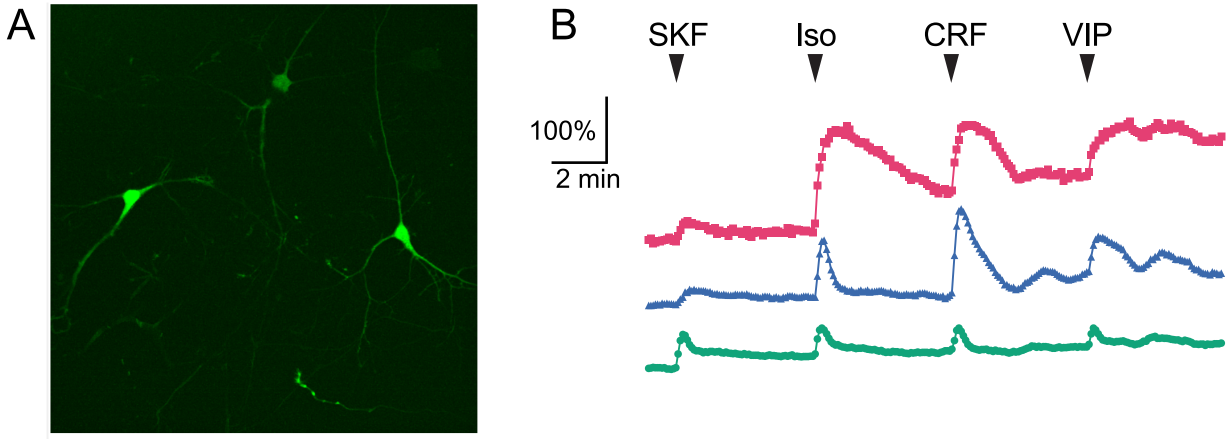


**Figure S1**
**A.** Image of 3 hippocampal neurons expressing cADDis. Scale bar, 100 µm. **B.** Fluorescent responses for each of the 3 neurons measured at the cell body, y axis is the % dF/F normalized to FSK application at the end (not shown). Sequential bath application of SKF (1 µM), Iso (1 µM), CRF (200 nM), and VIP (500 nM) is indicated by the arrows, with no washout between applications.

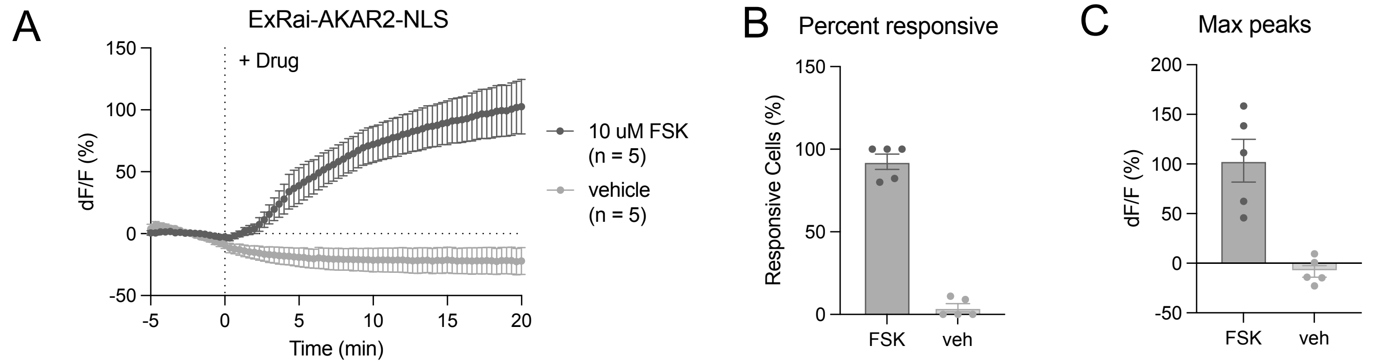


**Figure S2**

**A.** The change in fluorescence at the nucleus measured in neurons expressing ExRai-AKAR2-NLS upon addition of 10 uM forskolin (FSK) or vehicle. Images were taken every 20 sec. **B.** The percentage of imaged cells in a dish with a positive response, defined as reaching 20% change in fluorescence during the entire time course. **C.** The maximum fluorescence value from time point 0 – 20 min.

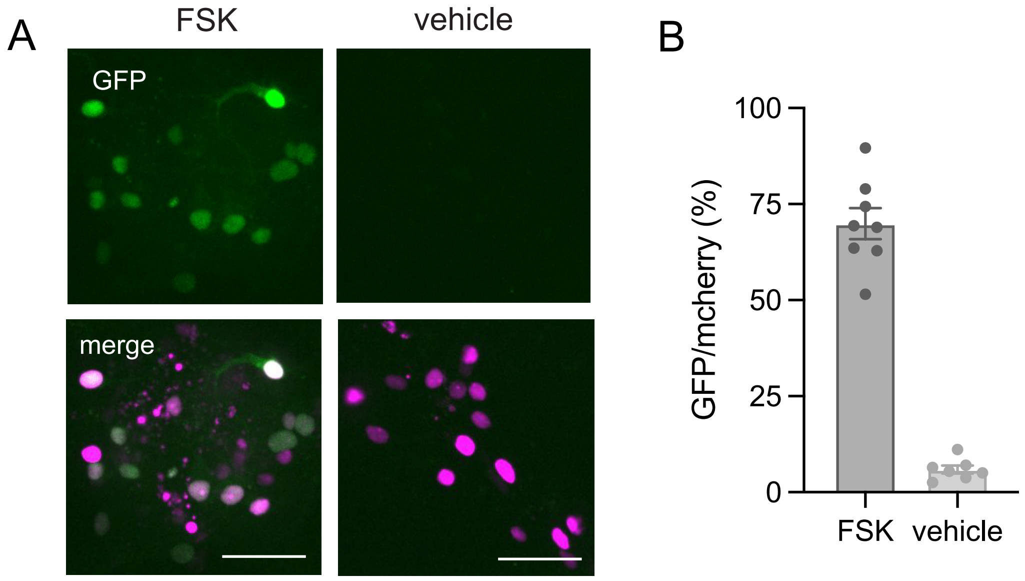


**Figure S3**
**A.** Representative images of hippocampal neurons expressing the CREB-GFP; UbC-mCherry reporter after 4 hours of incubation with TMP and 10 μM forskolin (FSK) versus vehicle. GFP marks cells where transcription was turned on, while mCherry (magenta) marks all cells that are expressing the reporter. Scale bar, 50 μm. **B.** The percentage of reporter-expressing neurons that have activated transcription after 4 hours of drug treatment. Each point represents the average of three 900 x 900 μm2 images from one dish from one neuron prep.

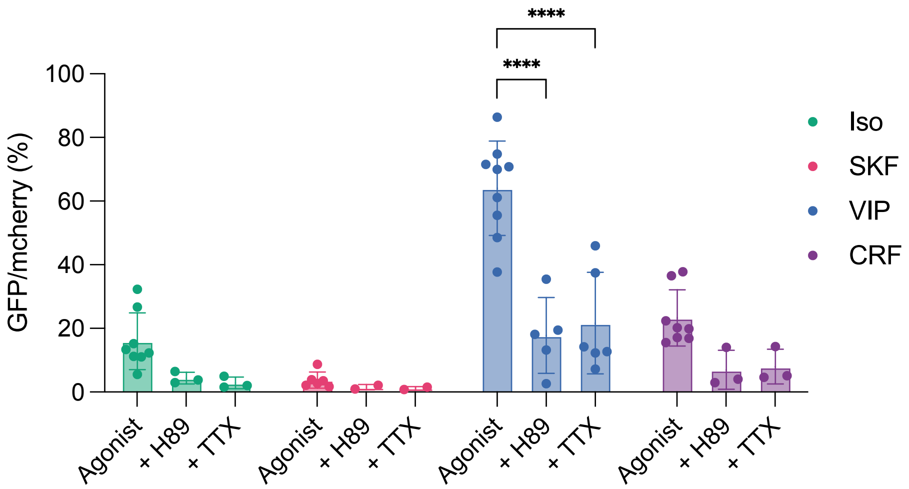


**Figure S4**
Quantification of the percentage of CREB-GFP; UbC-mCherry reporter-expressing neurons that have induced GFP expression after 4 hours of agonist treatment with and without inhibitors for PKA activity (H89, 20 μM) or sodium channels (tedrodotoxin, TTX, 500 nM). For inhibitor experiments, neurons were pretreated with H89 or TTX for 10 min prior to agonist addition. Two-way ANOVA with multiple comparisons (**** $p < 0.0001$).

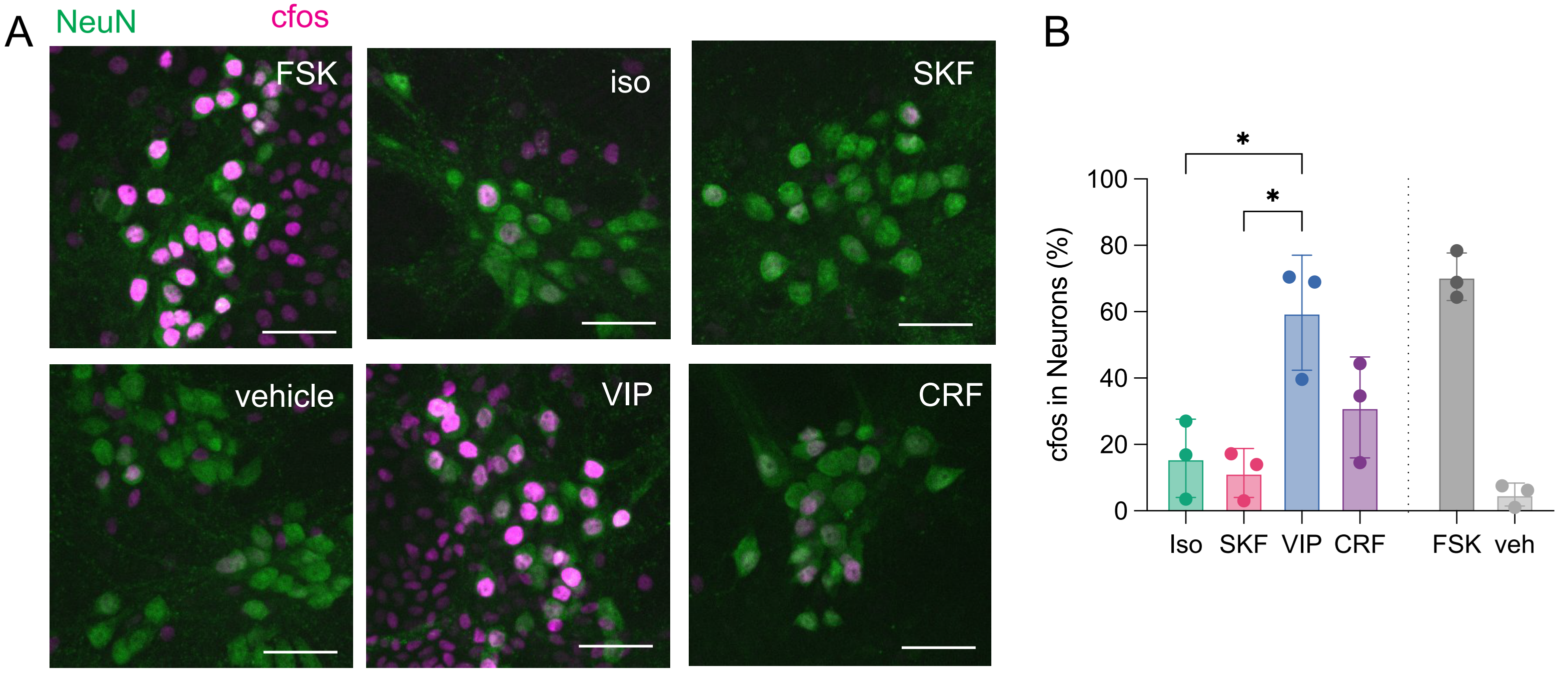


**Figure S5**
**A.** Representative images of hippocampal neurons that were treated with 1 hour of drug prior to fixation and immunostaining with antibodies for immediate early gene cfos (magenta) and neuronal marker NeuN (green). Scale bar, 50 μm. **B.** The percentage of neurons as marked by NeuN that have presence of cfos. Each point represents the average of three 900 x 900 μm2 images from one dish from one neuron prep. Ordinary one-way ANOVA with multiple comparisons (* $p < 0.05$).

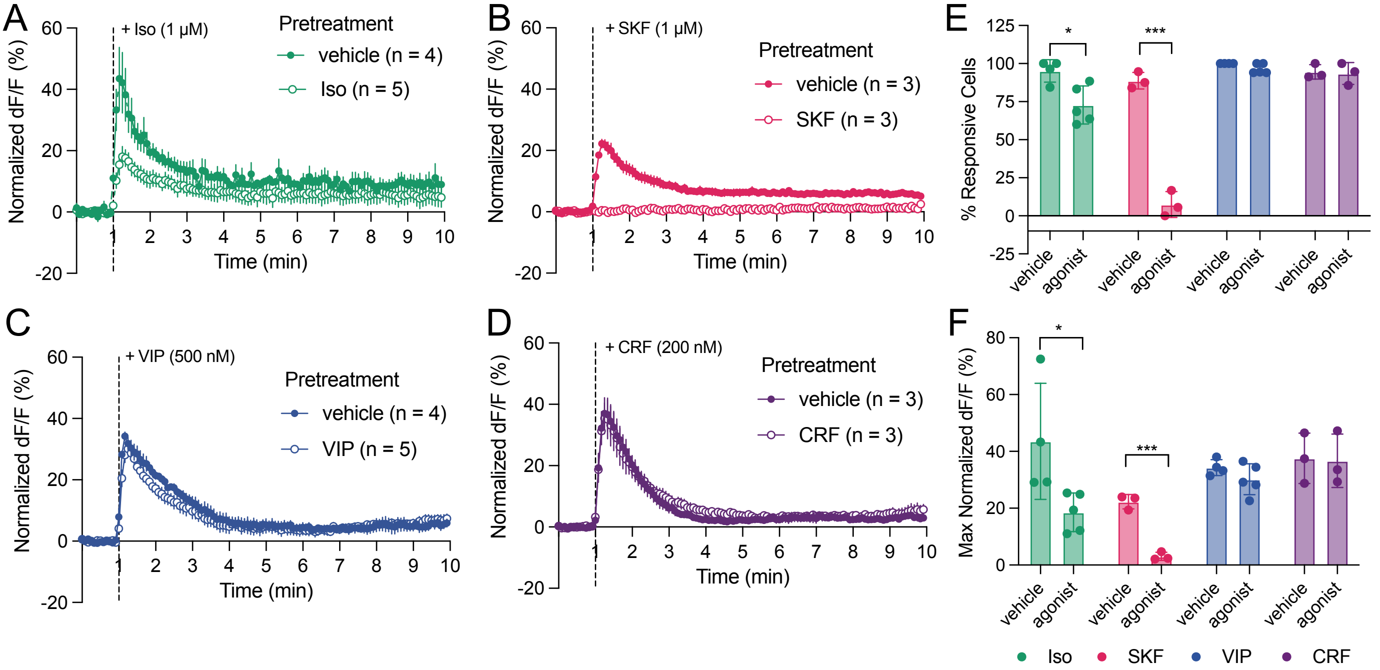


**Figure S6**

**A.** Neurons expressing the cAMP sensor cADDis were incubated with either 1 μM iso or vehicle for 10 minutes at 37 degrees. The dishes were then washed 3x with imaging solution and the fluorescence was measured across multiple cells upon addition of 1 μM Iso, normalized to 10 μM FSK. **B.** Normalized fluorescence changes upon application of SKF after 10 min pretreatment of SKF or vehicle. **C.** Normalized fluorescence changes upon application of VIP after 10 min pretreatment of VIP or vehicle. **D.** Normalized fluorescence changes upon application of CRF after 10 min pretreatment of CRF or vehicle. **E.** The percentage of cells in a dish that respond to the agonist following pretreatment by vehicle or the same agonist. Unpaired t-tests (* $p < 0.05$, *** $p < 0.005$). **F.** The maximum dF/F change of cells in a dish following pretreatment by vehicle or the same agonist. Unpaired t-tests (* $p < 0.05$, *** $p < 0.005$).
